# Optimized gene expression from bacterial chromosome by high-throughput integration and screening

**DOI:** 10.1101/2020.07.29.226290

**Authors:** Tatyana E. Saleski, Meng Ting Chung, David N. Carruthers, Azzaya Khasbaatar, Katsuo Kurabayashi, Xiaoxia Nina Lin

## Abstract

Chromosomal integration of recombinant genes is desirable compared to expression from plasmids due to increased stability, reduced cell-to-cell variability, and the elimination of antibiotics for plasmid maintenance. Here, we present a new approach for tuning pathway gene expression levels via random integrations and high-throughput screening. We demonstrate multiplexed gene integration and expression-level optimization for isobutanol production in *Escherichia coli*. The integrated strains could, with significantly lower expression levels than plasmid-based expression, produce high titers (10.0 ± 0.9 g/L isobutanol in 48 h) and yields (69 % of the theoretical maximum). Close examination of pathway expression in the top-performing, as well as other isolates, reveals the complexity of cellular metabolism and regulation, underscoring the need for precise optimization while integrating pathway genes into the chromosome. This new method for multiplexed pathway gene integration and expression optimization could be readily extended to a wide range of pathways and chassis to create robust and efficient production strains.

## Introduction

Microbial biosynthesis is a sustainable, high-specificity approach to achieving chemical conversions with the potential to produce a vast assortment of pharmaceutical, fuel, and commodity marginchemicals. Development of a high-performing production strain for a desired molecule generally requires tuning the expression levels of native and/or heterologous genes in hosts such as *Escherichia coli*^1, 2^. While extrachromosomal multicopy plasmids offer a convenient method for rapidly prototyping different expression levels of genes, and the majority of metabolic engineering efforts utilize them, they have several challenging attributes. First and foremost, plasmids are unstable, which has been extensively documented and studied due to the importance of plasmids in biotechnology^3–9^. Cells with plasmids can suffer from both structural instability, in which the plasmid is still carried but mutations inactivate the gene of interest, as well as segregational instability, in which some cells no longer carry the plasmid^10, 11^. Plasmids can also multimerize, which decreases their stability^12, 13^. Second, plasmid copy number can vary between cells within stable populations, and the degree of variation is often not well characterized even for widely used plasmids^14^. Münch *et al.* presented a striking example of cellular heterogeneity during plasmid-based production of recombinant protein in *Bacillus megaterium*. The authors found that, under the condition of strong selective pressure, 30% of the cell population was in a low-producing state^15^. This heterogeneity was found to be a result of asymmetric plasmid distribution. Moreover, plasmids require a selective pressure, which is typically an antibiotic molecule. The addition of antibiotics raises the process costs and puts additional stress on the cells. It is also not always effective. Kanamycin, for example, can become an ineffective selection agent at high phosphate concentrations^16^. Furthermore, due to the rise of multidrug-resistant pathogens, there is strong motivation to reduce antibiotic usage^17^.

Chromosomal integration and expression of genes is an effective way for circumventing the issues of plasmids described above. This alternative approach leads to more stable and robust production strains, as well as reduce leakiness of inducible expression systems, which are highly desirable traits for large-scale and long-term production processes^18, 19^. Mairhofer *et al.* compared a plasmid-based to a chromosomally integrated expression system and found a much more significant stress response in the plasmid-carrying strain. They identified massive overtranscription of the plasmid-based gene of interest, leading to diversion of ribosomes and other cellular resources, as the source of the metabolic burden^20^.

Significant research efforts have been devoted to developing techniques for synthetic integrations^21–23^. The two tools most widely used in prokaryotes for targeted integration of constructs are homologous recombination by expression of the RecET or λ-Red proteins^24–26^ and site-specific recombination^27^. Homologous recombination enables rapid integration into a desired locus but is only suitable for constructs up to several kilobases in size and that do not carry significant homology to another part of the genome. Site-specific recombination is less restrictive of the input construct but requires a bacterial chromosomal attachment site (*attB*) in the recipient genome, which can place limitations on the location of integration. This site restrictiveness can be circumvented by first inserting the attachment site into the target locus^28^. Yet, despite the recognized desirability of chromosomal integration and the genetic tools available for targeted integrations, achieving suitable gene expression levels from the chromosome for optimal production remains far more challenging than manipulation of gene expression via plasmids, even in well-studied organisms such as *E. coli*^22^.

Chromosomal gene expression levels are dependent on multiple determinants. Similar to plasmid-based gene expression, elements of the genetic construct, including promoters, ribosome-binding sites (RBSs), and enhancers or activators, can be exploited to optimize heterologous gene expression. Yet, single-copy expression from the chromosome is generally weaker than that of the same construct from a multi-copy plasmid. Subsequently, there have been efforts in increasing the copy number on the chromosome to achieve high-level gene expression^18, 29^.

Another determinant affecting the expression level from the chromosome strongly is the location of gene integration. It has long been known that during exponential growth, genes closer to the origin of replication experience higher gene dosage and, therefore, higher expression levels^30, 31^. More recently it has been shown that other factors of genomic position, such as level of DNA compaction or proximity to active genes, can also affect gene expression levels^32, 33^. In examining transcription levels of a reporter gene at various sites in the *E. coli* genome, Bryant *et al.*^32^ and Scholz *et al.*^33^ observed significant differences, up to ~300-fold, in expression across the *E. coli* genome, excluding gene dosage effects. Intriguingly, on the other hand, it has been suggested that in nature, microorganisms may use transposon-mediated transfer of catabolic genes to different genomic locations as a means of fine-tuning their expression levels in order to adapt to new environmental conditions^34^. These findings suggest that modulation of gene integration position can serve as a useful tool for regulating expression levels. For instance, Loeschcke *et al.* have developed an approach for transfer and expression (TREX) of biosynthetic pathways in bacteria utilizing Tn5 transposase to integrate gene clusters into random sites of the chromosomes and, based on resulting colony color intensities, observed a range of production levels for pigmented secondary metabolites^35^.

Synthetic biology applications targeting the production of a small molecule often require delicately balanced cellular resource allocation between production and growth, as well as across different genes of the biosynthetic pathway. It remains elusive to predict the optimal gene expression levels for most pathways due to the highly entangled nature of cellular metabolic and regulatory networks. Additionally, for chromosomal gene expression, it is not fully understood yet how the expression of different regions of the chromosome is affected by changes in environmental conditions and whether/how an inserted gene construct will affect the native expression level of the insertion region. It is, therefore, very difficult to rationally decide where to integrate a gene construct. Here, we present a new metabolic engineering strategy for optimizing a pathway’s performance by tuning the gene expression through position-dependent expression variation (Fig. 1). In this method, we utilize the Tn5 transposase to randomly integrate pathway genes in the *E. coli* genome in a multiplexed fashion. We subsequently screen the libraries using syntrophic co-culture amplification of production (SnoCAP), which converts the production phenotype into a readily screenable growth phenotype, as described in our previous work^36^. We demonstrate the approach using a pathway for the production of isobutanol, a promising drop-in biofuel^37^.

**Figure 1.**
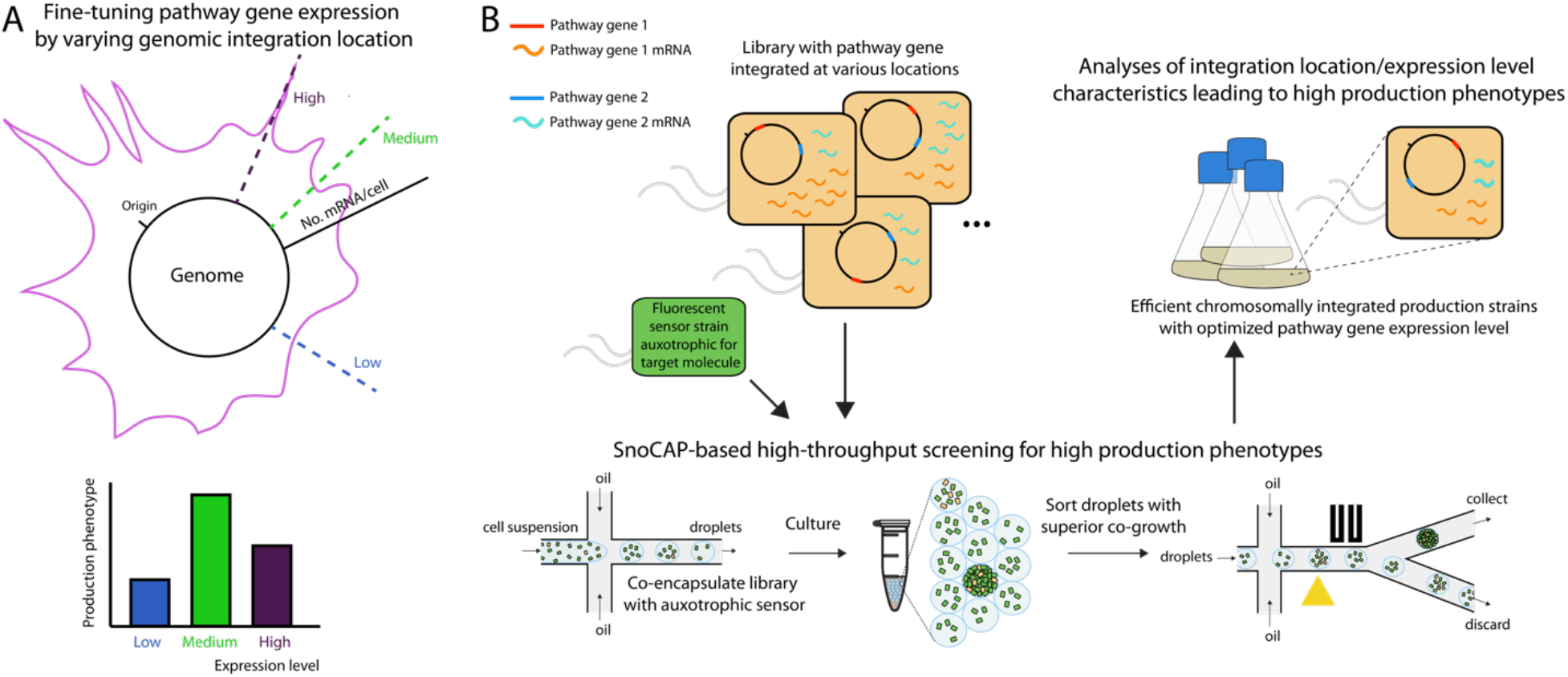
New approach for optimizing chromosomal heterologous gene expression by position-dependent expression variation. (a) Gene expression level varies depending on distance from the origin of replication, as well as other position-dependent factors. The expression level of a heterologous pathway gene affects the production phenotype of a strain. Increasing the expression level of a gene can improve production level by increasing flux through the pathway, but avoiding too high of an expression level can also provide benefits such as reducing the buildup of toxic intermediates and avoiding unnecessary metabolic burden. (b) Libraries with pathway genes integrated into various genome locations can be screened for production phenotype using syntrophic co-culture amplification of production (SnoCAP)^36^. A cross-feeding co-culture is configured consisting of a secretor strain producing a target molecule and a sensor strain that cannot synthesize the target molecule autonomously. The co-cultures are compartmentalized in microfluidic droplets, incubated to allow co-growth, and then sorted to identify the strains with high production phenotypes.

Efficient isobutanol production in *E. coli* has been achieved by diverting the branched-chain amino acid pathway intermediate 2-ketoisovalerate (2-KIV) for conversion into isobutanol^38^ (Fig. 2A). It has been demonstrated that by overexpressing AlsS from *Bacillus subtilis*, IlvC and IlvD from *E. coli* (for the conversion of pyruvate to 2-KIV), and KivD and AdhA from *Lactococcus lactis* (for the conversion of 2-KIV to isobutanol), production can reach up to 84% of the theoretical yield under aerobic batch conditions^38^ and titers can reach up to 50.8 g/L under fed-batch conditions^39^. Resolution of a cofactor imbalance has enabled strains to reach 100% of theoretical yield under anaerobic conditions^40^, and a recently developed strategy for isobutanol-linked growth selection under anaerobic conditions enabled optimization of pathway expression levels for achieving optimal production rates^41^.

**Figure 2.**
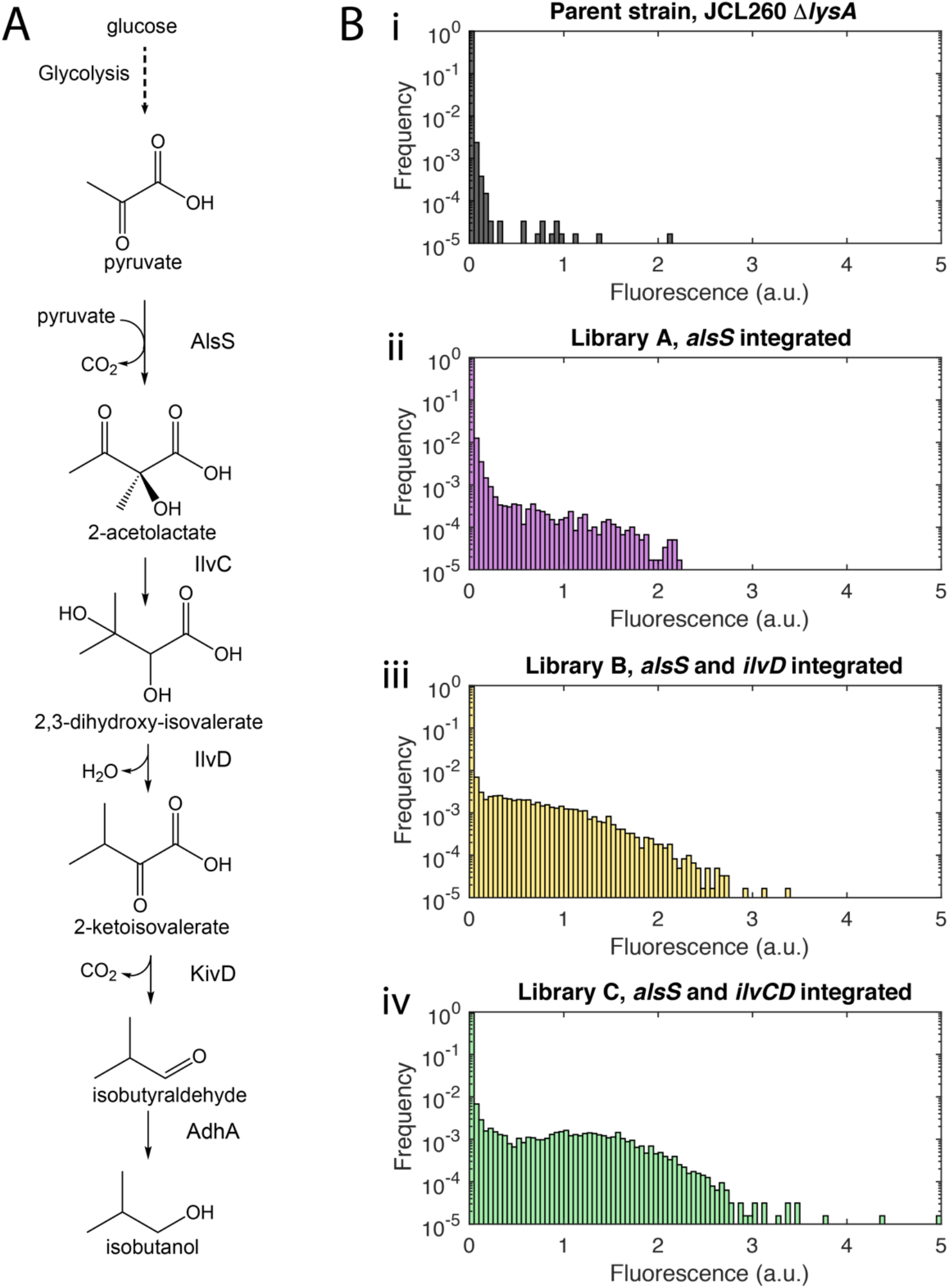
Screening of libraries with *alsS*, *ilvCD* integrated by Tn5. (a) Isobutanol production pathway^50^. Acetolactate synthase (AlsS), ketol-acid reductoisomerase (IlvC), dihydroxy-acid dehydratase (IlvD), 2-ketoacid decarboxylase (KivD), and alcohol dehydrogenase (AdhA). (b) Histograms comparing fluorescence signal from droplets containing co-cultures of fluorescent sensor strain with unlabeled secretor strain libraries, compared to the parent strain (i). The libraries were constructed by Tn5 integration of *kan-*PLlacO1*-alsS* (Library A) (ii), PLlacO1-*ilvD* followed by *kan-*PLlacO1*-alsS* (Library B) (iii), or PLlacO1-*ilvCD* followed by *kan-*PLlacO1*-alsS* (Library C) (iv). Each histogram comprises data from the analysis of about 60,000 droplets.

While much work with this pathway has relied on plasmid-based expression of pathway genes, there is also interest in developing chromosomally integrated strains. Akita et al. have demonstrated the integration of the five pathway genes into the *E. coli* genome, resulting in a titer of 6.8 g/L and a yield of 55%, without the addition of costly antibiotics or inducers^42^. Bassalo et al. developed a new CRISPR-based method for high-efficiency, single-step chromosomal

In the present work, we demonstrate that the generation of libraries with isobutanol pathway genes integrated into diverse chromosomal positions combined with high-throughput production phenotype screening is an effective means of generating high-performance chromosomally integrated strains. *E. coli* strains resulted from this study show, to our knowledge, the highest titers and yields yet reported with the isobutanol pathway genes expressed exclusively from the chromosome.

## Results

### Construction and screening of Tn5 integration libraries

We began by constructing an integration library (Library A) in which a construct containing the *alsS* gene, under the PLlacO1 promoter, and a kanamycin-resistance gene, was integrated by Tn5 transposase into the genome of JCL260 Δ*lysA* in random locations. This integration was accomplished by PCR amplification of the *alsS* and *kan* genes with primers that add the 19 bp mosaic end sequence recognized by Tn5, followed by reaction with Tn5, and electroporation of cells with the resulting transposome. The approximate library size was determined to be ~3.4 ± 1.1 × 10^4^ by plating a small quantity of the cells after a short initial recovery on kanamycin-containing plates. The remainder of the library was subjected to a liquid kanamycin selection and subsequently screened by SnoCAP. In the SnoCAP approach, the library is co-encapsulated in water-in-oil microdroplets with a fluorescent sensor strain that is auxotrophic for the target molecule of interest. The library itself is also auxotrophic for a second, orthogonal molecule supplied by the sensor strain. This cross-feeding configuration enables conversion of the production phenotype into a fluorescent output, amplified by co-culture growth^36^. To screen for overproducers of 2-ketoisovalerate, an intermediate in the isobutanol pathway (Fig. 2A), we use a sensor strain with a deletion of the *ilvD* gene (encoding a dihydroxy-acid dehydratase), which prevents it from converting 2,3-dihydroxy-isovalerate into 2-ketoisovalerate. This strain can only grow when it is provided an exogenous source of 2-ketoisovalerate and/or branched-chain amino acids. We demonstrated in our previous work that this sensor can be used to identify strains that overproduce isobutanol once the genes responsible for converting 2-ketoisovalerate to isobutanol (*kivD* and *adhA*) are added^36^.

The cells are loaded at densities such that all droplets contain sensor cells, and most droplets contain either zero or one secretor cell from the integration library. Thus, the co-growth phenotype of individual library members may be assessed. The microdroplet compartmentalization format enables high-throughput screening and isolation of droplets of interest by fluorescence-activated droplet sorting (FADS)^43^.

Fluorescence measurement of the droplets containing Library A members co-cultured with fluorescent sensor strain showed that more droplets exhibited growth than when droplets contained co-cultures with the parent strain, which lacks the *alsS* gene, as the secretor (Fig. 2Bi,ii). We sorted the top 0.3% of droplets with the highest fluorescence signal and plated the pool of droplets on plates with tetracycline to select for the secretor cells and against the sensor cells (the parent strain of the secretor strain libraries contains a tetracycline resistance gene in the F plasmid, whereas the sensor strain is tetracycline sensitive). We assessed the isobutanol production of colonies obtained from the screening and colonies from the unscreened library, after transformation with pSA65. Of 12 colonies chosen at random from the unscreened library, 10 produced less than 0.1 g/L after 48 h of fermentation (similar to the parent strain), one produced 0.4 g/L and the other produced 2.1 g/L (Fig. S1). Of 10 random colonies tested from the sorted pool, all produced more than 2.5 g/L (Fig. S1), which demonstrates the effectiveness of the screening method.

Because transposome integration efficiency is expected to decrease with increasing construct length, we chose to use the *ilvCD* genes as a selection marker by utilizing their essentiality in minimal medium, rather than an antibiotic. We deleted *ilvD* from JCL260 Δ*lysA* and integrated constructs containing either both genes *ilvC* and *ilvD* behind the PLlacO1 promoter or only the *ilvD* gene behind the same promoter. We performed the selection in minimal M9 medium supplemented with IPTG and lysine, with a small amount plated on solid medium for library size estimation and the rest of the library undergoing liquid selection. These integrations produced far smaller library sizes than the *alsS* integration construct (~10^2^ integrants per electroporation vs. ~10^4^ for *alsS*). We determined that this was due to substantially lower survival rates in the minimal medium selection compared to kanamycin selection in LB medium. For example, when JCL260 Δ*lysA* Δ*ilvD* was transformed with pSA69, the number of colonies recovered on LB plates with kanamycin was two orders of magnitude higher than the number on M9 plates with lysine and IPTG. Further optimization of the recovery method, such as supplementation of additional nutrients or seeding of small quantities of the branched-chain amino acids may help improve this lower-than-expected recovery rate and increase the diversity of the library. It should also be noted that the survival rate on the solid medium may be different from that in the liquid medium, which could contribute to inaccuracies in estimated library size. Deep sequencing of the library could be used to quantify the library size more accurately. We repeated the *ilvD* and *ilvCD* library generation several times and pooled the integrants, resulting in libraries of each of size ~500-1,000 (estimated from agar plating). We then integrated the *alsS-kan* transposome into these libraries, which led to ~1.1 ± 0.5 × 10^4^ integrants for the *ilvD/alsS* library (Library B) and 1.3 ± 0.3 × 10^4^ for the *ilvCD/alsS* library (Library C). Therefore, the library should consist of about 10 different *alsS* integration locations for each *ilvD* or *ilvCD* integration library member.

When assessed by the SnoCAP method (library members co-encapsulated in microdroplets with sensor strain, incubated for co-growth, and the fluorescence of the droplets measured), the *ilvD* and *ilvCD* integration libraries displayed no measurable co-growth. The *ilvD/alsS* and *ilvCD/alsS* libraries (Libraries B and C, respectively) on the other hand, showed substantially more fluorescence compared to the library with only *alsS* integrated (Library A) (Fig. 2B).

We sorted the droplets containing co-cultures of these libraries and recovered colonies from the pools of droplets, and then transformed the isolates with plasmid pSA65 (encoding KivD and AdhA) and assessed isobutanol production. The best isolates were found to be from Library C. In particular, an isolate termed C7 reached a production level closely approaching that of the strain expressing the pathway genes from a plasmid (Fig. 3A). With the goal of understanding the relationship between production phenotype and genome-position-dependent gene expression, we decided to examine isolates displaying a spectrum of production levels and hence also isolated several library members with low- and intermediate-production levels (Fig. 3A) by screening the random libraries via the microplate format of SnoCAP.

**Figure 3.**
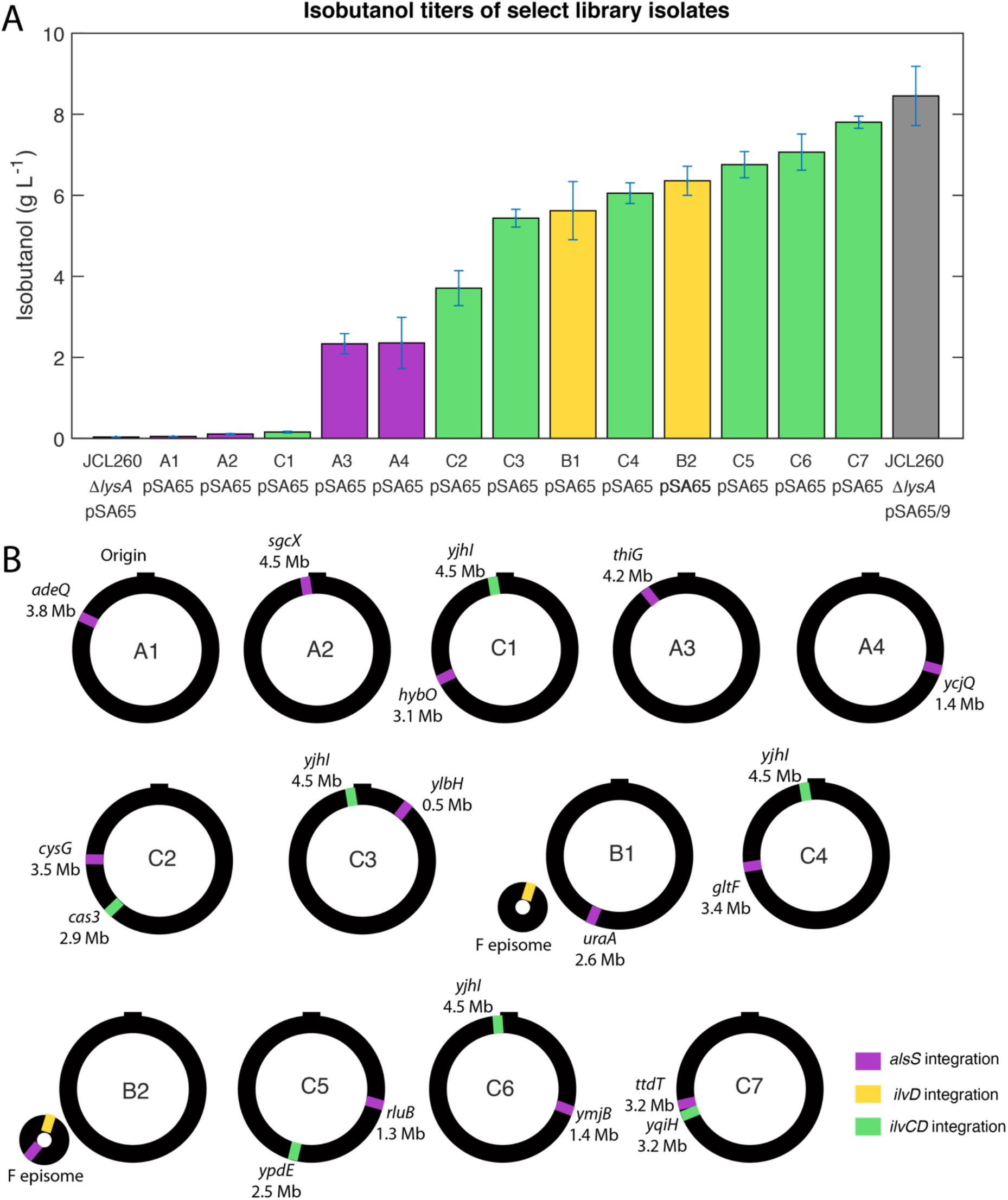
Production phenotype and gene insertion locations of select library isolates. a) Isobutanol titers of library isolates displaying various production levels. Samples were taken 48 h after induction of the pathway genes. Each of the isolates has been transformed with the pSA65 plasmid expressing *kivD* and *adhA*. Parent strain JCL260 Δ*lysA* and JCL260 Δ*lysA* pSA69, which expresses the *alsS* and *ilvCD* genes from the pSA69 plasmid, each also carrying pSA65, are included for comparison. Error bars represent the standard deviation of three biological replicates. (b) Gene integration locations of the isolates whose production is shown in (a).

### Genomic characterization of select library isolates

We then determined the insertion site locations in the selected isolates by transposon footprinting, which also allowed us to eliminate duplicates of the same genotype and resulted in a panel of unique isolates reported in this work. The integration locations were found to spread across the whole genome (Fig. 3B, Table 1). Intriguingly, each of the Library B isolates, B1 and B2, contains integrations into the F plasmid, a large (~100 kb) plasmid previously introduced to the base strain to supply *lacIq* for increased levels of the lac repressor. The F plasmid is stably maintained in *E. coli* due to an active partitioning system^44, 45^, and it has been suggested that placing heterologous genes on the F plasmid may provide improved stability over the use of other plasmids^46, 47^. It is also interesting that in isolate B2, both integration constructs are found to be located on the F plasmid.

**Table 1.** Gene integration locations.

| Strain | Integration construct | Gene | Coordinate in gene of integration location | Direction relative to gene integrated into |
| --- | --- | --- | --- | --- |
| A1 | *kan-alsS* | *adeQ* | 436 | Opposite |
| A2 | *kan-alsS* | *sgcX* | 111 | Opposite |
| C1 | *kan-alsS* | *hybO* | 812 | Opposite |
| C1 | *ilvCD* | *yjhl* | 231 | Same |
| A3 | *kan-alsS* | *thiG* | 73 | Same |
| A4 | *kan-alsS* | *ycjQ* | 364 | Opposite |
| C2 | *kan-alsS* | *cysG* | 563 | Opposite |
| C2 | *ilvCD* | *cas3* | 1,387 | Same |
| C3 | *kan-alsS* | *ylbH* | 205 | Opposite |
| C3 | *ilvCD* | *yjhl* | 231 | Same |
| B1 | *kan-alsS* | *uraA* | 659 | Same |
| B1 | *ilvD* | gamma delta (Tn1000) region of F plasmid | 426 bp into *tnpX* | Same as *tnpX* |
| C4 | *kan-alsS* | *gltF* | 313 | Opposite |
| C4 | *ilvCD* | *yjhl* | 231 | Same |
| B2 | *kan-alsS* | Upstream of *psiB* on F plasmid | 218 bp upstream | Opposite of *psiB* |
| B2 | *ilvD* | gamma delta (Tn1000) region of F plasmid | 426 bp into *tnpX* | Same as *tnpX* |
| C5 | *kan-alsS* | *rluB* | 794 | Same |
| C5 | *ilvCD* | *ypdE* | 76 | Same |
| C6 | *kan-alsS* | *ymjB* | 7 | Opposite |
| C6 | *ilvCD* | *yjhl* | 231 | Same |
| C7 | *kan-alsS* | *ttdT* | 114 | Opposite |
| C7 | *ilvCD* | *yqiH* | 512 | Opposite |

Another noteworthy observation was that two of the disrupted genes, *ttdT* and *yjhI*, are genes in which we had found mutations during previous work in which we chemically mutagenized JCL260 Δ*lysA*, screened for improved isobutanol productivity and resequenced the genome of an improved isolate^36^. In that study, the strain with the best production level possessed a mutation in the *ttdT* gene leading to an alanine to valine substitution at position 390 of the protein, and *yjhI* contained a mutation leading to a premature stop codon after the first 187 amino acids (of 227).

In the best isolate, C7, the two integration constructs landed in *ttdT* and *yqiH,* two genes that are quite close to each other, separated by only 18 kb (Fig. 4A). To understand the genetic mechanism underlying the high-production phenotype of this isolate, we investigated whether the gene disruptions, besides expression of the inserted genes, can contribute to increased production. We knocked out the *ttdT* and *yqiH* genes individually in a base strain JCL260 Δ*lysA*.

**Figure 4.**
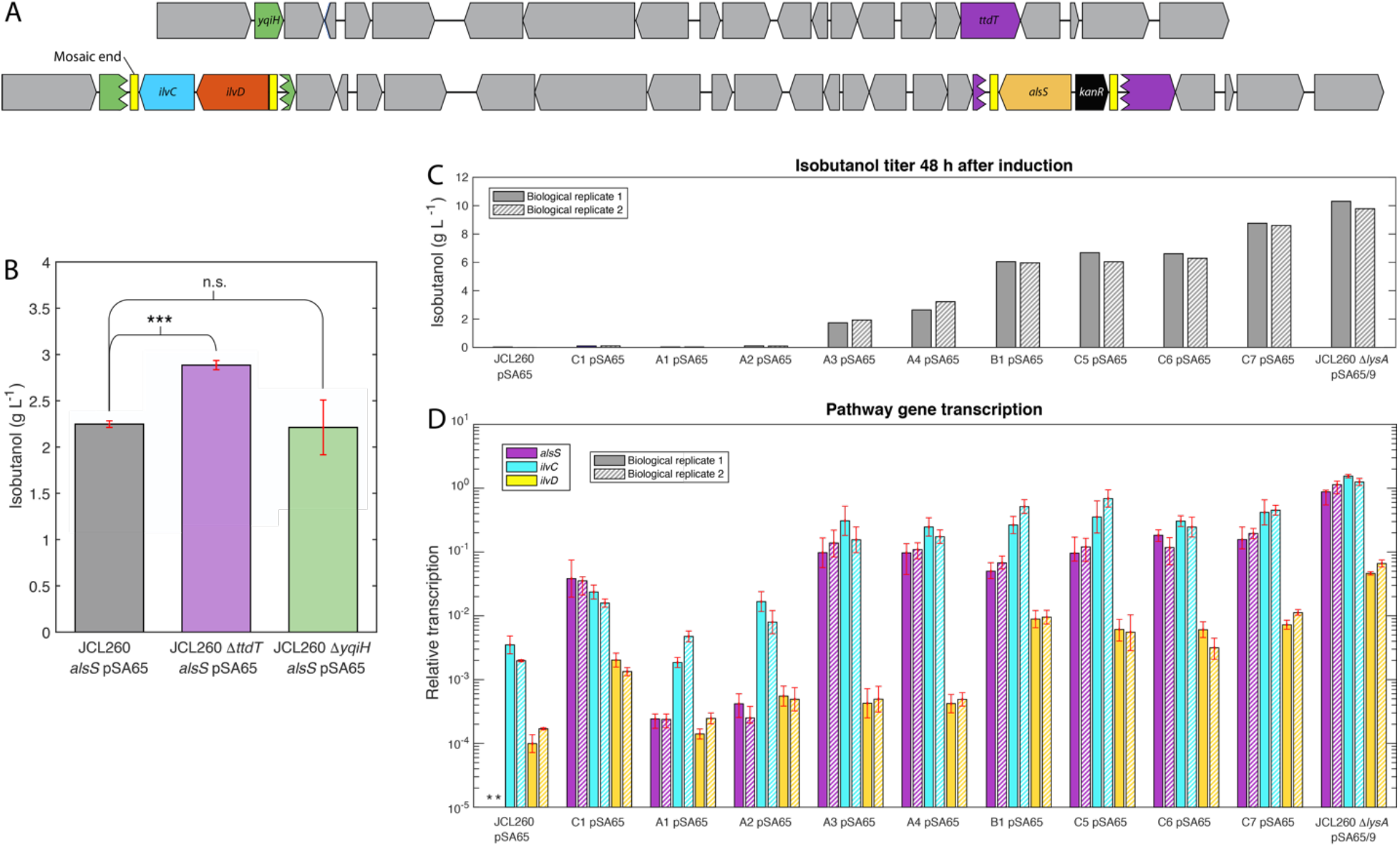
Investigation of mechanisms underlying production phenotypes of select library isolates. Schematic of the locations of *kan-*PLlacO1*-alsS* and PLlacO1-*ilvCD* integrations in isolate C7, determined by transposon footprinting. (B) Isobutanol production of strains with *alsS* genomically integrated, containing plasmid pSA65 encoding *kivD* and *adhA*, and possessing deletions of either *ttdT* or *yqiH*, compared to the same strain without either deletion. Data shown are from samples taken 72 h after induction with IPTG. ***: significant difference at *P*-value < 0.0001 using an unpaired t-test. n.s.: not significant. (C) Isobutanol production 48 h after induction of the cultures from which the RNA had been harvested. (D) Expression levels of *alsS*, *ilvC*, and *ilvD*, relative to *alsS* level in JCL260 Δ*lysA* pSA65/9. RNA was harvested 3 h after induction with IPTG. * indicates no amplification during the qPCR reaction. Data from two biological replicate cultures of each strain are shown. Error bars in (B) represent the standard deviation of three biological replicates. Error bars in (D) represent the standard deviation of three technical replicates.

Into each deletion strain, we then integrated *alsS* in the *yghX* site, which we have previously seen leads to an intermediate production level (and therefore should allow us to see changes in the production that may not be visible in a very low- or high-producing background strain). After transformation of the strains with pSA65, we tested isobutanol production and found that the *ttdT* deletion strain produces more isobutanol to a statistically significant degree compared to the strain possessing wildtype *ttdT*, whereas *yqiH* deletion does not have a statistically significant effect (Fig. 4B).

### Characterization of pathway gene transcript levels in select library isolates

We next analyzed gene expression in various isolates of different production levels. We harvested the RNA during exponential growth (3 h after induction of the pathway genes with IPTG), reverse transcribed it and analyzed gene expression of *alsS*, *ilvC*, and *ilvD* by quantitative real-time PCR (qPCR) (Fig. 4D), while meanwhile continuing to grow the cultures and measuring their isobutanol production 48 h following induction (Fig. 4C). It was found that the pathway gene expression in each of the integrated strains was, as expected, far lower than that in the strain expressing *alsS/ilvCD* from the pSA69 plasmid (Fig. 4D, note the log-scale of the Y-axis). Yet, with significantly lower expression levels of all three genes, the top isolate C7 showed a production level very close to that of the plasmid strain.

With the gene transcript profiles of these isolates, we sought to elucidate the relationship between gene expression and production phenotype. Some interesting observations were made. For instance, predictably, isolates from Library A have lower expression of *ilvD* than isolates from Libraries B and C. However, *ilvC* can be seen to be upregulated even in certain strains that do not have *ilvC* insertion (A3, A4, B1). These strains have high expression levels of *alsS*, so it is likely that accumulation of acetolactate, the product of AlsS, is inducing higher transcription levels of *ilvC* in these strains, as has been previously observed^48^. We also noted that there was a similar trend of relative expression levels within the pathway (*ilvC* highest followed by *alsS* and then *ilvD*) in the medium- and high-production isolates (A3, A4, B1, C5, C6, C7), which was also seen in the plasmid strain. Most strikingly, however, there was a lack of a clear relationship between gene expression levels and production. In particular, despite the fact that the top four analyzed isolates (B1, C5, C6, C7) all showed a much higher expression of *ilvD* than the next two, substantially lower-producing, isolates (A3, A4), it is not clear what causes C7 to perform significantly better in production than the other three isolates or why differences in gene expression levels across these three do not lead to differences in production. Examination of the low-production isolates also provided some insights. Low expression, especially of *alsS*, appeared to have caused low production in A1 and A2. Intriguingly, another isolate C1 showed much higher expression of the three pathway genes but did not exhibit increased production. We suspect that this could be due to deleterious effects of disrupting the genes in the integration sites or to poor pathway balancing. The above findings underscore the highly entangled and complex nature of optimizing pathway gene expression for a specific production phenotype.

### Construction and characterization of strains with all isobutanol pathway genes integrated

To generate strains in which the entire isobutanol pathway is expressed from the chromosome, we integrated the *adhA* and *kivD* genes into isolate C7. Because in the work of Atsumi et al.^38^, these genes were expressed from a high-copy-number plasmid, we expected that high expression levels would be needed to achieve suitable production levels. We utilized the chemically inducible chromosomal evolution (CIChE) method of Tyo et al.^18^ for generating multiple copies of a gene in a single integration site. In this approach, the genes of interest are placed alongside an antibiotic marker between 1 kb homology regions that enables RecA-mediated recombination to increase the copy number in response to increases in antibiotic concentration (Fig. 5A). If desired, the copy number can be stabilized by deletion of the *recA* gene. We integrated the CIChE construct containing *kivD, adhA,* and the chloramphenicol resistance gene *cat* into the *aslB* site, a high-expression site^32^. Before integrating this construct into C7, we also integrated it into a clean background strain, passaged it into higher chloramphenicol concentrations, deleted the *recA* gene to stabilize the copy number, and measured the construct copy number by qPCR of the *cat* gene (Fig. S2). These measurements demonstrated that the copy number can be increased up to about 60 copies by this method. We tested the production of the C7 strain with the CIChE *kivD-adhA* integration (C7 CIChE) in 50 μg/mL chloramphenicol and found that it produced more isobutanol than a strain with just a single copy of *kivD* and *adhA* integrated into the same site. Further increasing the chloramphenicol concentration did not lead to an increase in production level. The fully integrated strain C7 CIChE performed very similarly to C7 pSA65, generating isobutanol titers and yields lower than but close to those of the double plasmid strain (Fig. 5B,C). Noticeably, on the other hand, this strain reached higher cell densities than the plasmid-bearing strains (Fig. 5D).

**Figure 5.**
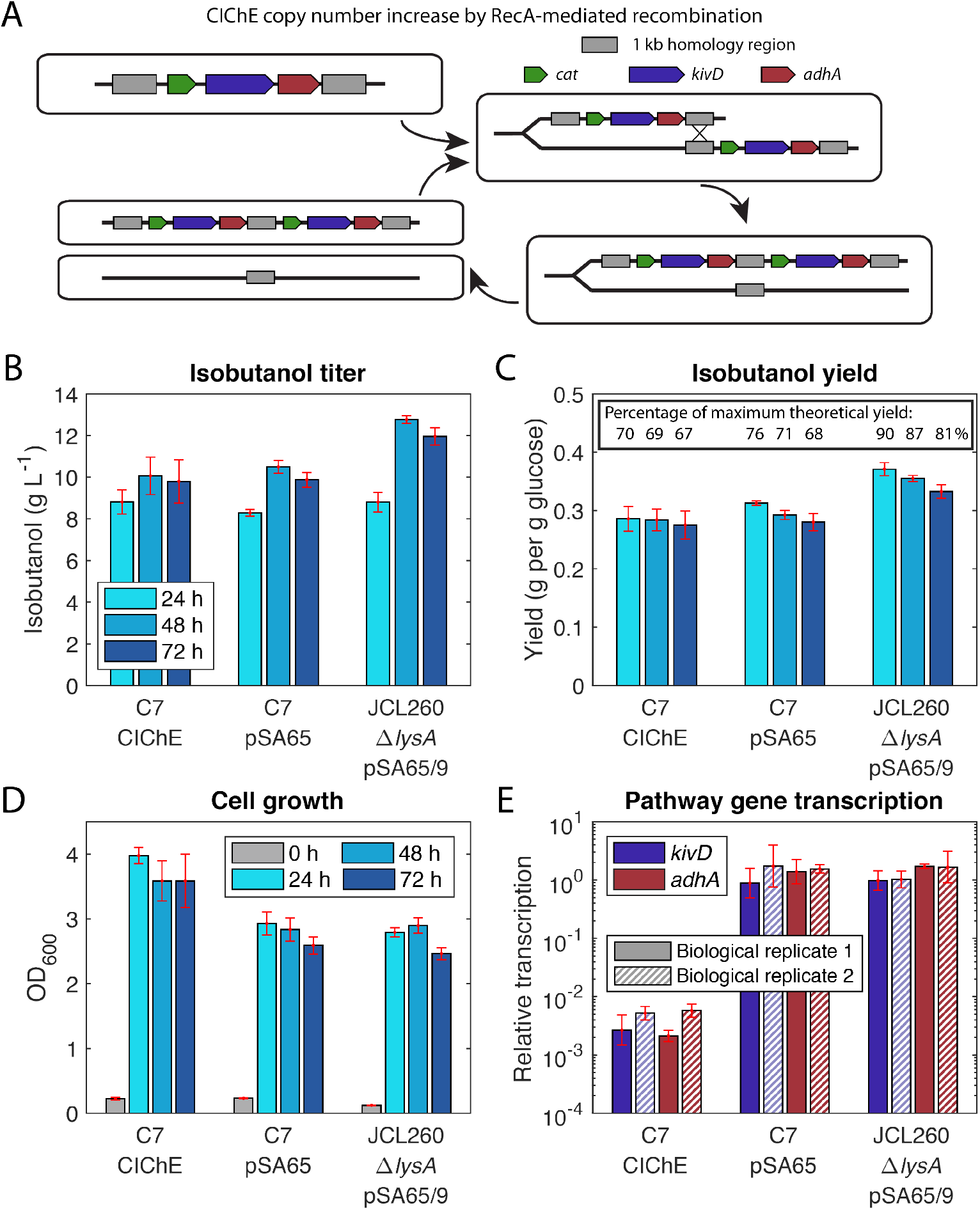
Fully integrated strain performance compared to strains with plasmids. (a) Overview of the chemically inducible chromosomal evolution (CIChE) approach for gene copy number amplification by RecA-mediated homologous recombination. (b) Isobutanol production by strain C7 after integration of *kivD-adhA* by CIChE, compared to C7 pSA65, which has *alsS* and *ilvCD* integrated into the genome and *kivD* and *adhA* expressed from the pSA65 plasmid, and JCL260 Δ*lysA* pSA65/9 which has all five pathway genes expressed from the plasmids. Error bars represent the standard deviation of three biological replicates. (c) The yields of the cultures based on glucose consumed. The inset in (c) lists the percentage of the theoretical maximum yield (0.41 g isobutanol per gram glucose). (d) The cell densities of the production cultures whose performance is displayed in (b) and (c). (e) Expression levels of *kivD* and *adhA*, relative to *kivD* level in JCL260 Δ*lysA* pSA65/9. RNA was harvested 4 h after induction with IPTG. Error bars represent the standard deviation of three technical replicates.

We also examined the *kivD* and *adhA* transcript levels in the fully integrated strain compared to the plasmid-bearing strains. As expected, they are lower in C7 CIChE. Quite surprisingly, however, we found them to be dramatically lower (more than two orders of magnitude) in the integrated strain compared to those in both strains expressing *kivD*/*adhA* from plasmid pSA65 (Fig. 5E). This indicates that the expression levels rendered by the high-copy number plasmid are unnecessarily high. This result further demonstrates that it is feasible to achieve high production through significantly lower pathway gene expression from the chromosome.

## Discussion

Synthetic biology is making strides toward solutions to some of society’s most pressing problems. Managing the issue of stability and cell-to-cell variability, however, still present major hurdles to overcome^49^. Moving gene expression from plasmids to the genome can improve stability and reduce cell-to-cell variability. Yet it remains challenging to predict the expression levels that will be optimal for production and achievable by genomic integration into a particular site. Moreover, the interplay between gene integration and surrounding regions on the chromosome, such as gene disruption and expression level perturbation, could cause complications and additional difficulty in optimizing pathway expression.

The new approach of screening of random integration libraries developed in the present study circumvents the challenges of rational selection of integration locations. The transposon integration method is versatile and applicable to almost any gene of interest. Since the integration is not homology-based (unlike lambda Red recombinase-based integration, for example) it can even be used on genes that are already natively present in the genome at suboptimal expression levels, such as *ilvC* in the work presented here. The high-throughput screening enables large libraries to be assessed, making it feasible to multiplex integrations of different pathway genes.

The application of the approach to the isobutanol pathway demonstrates its effectiveness and has generated a strain with, to our knowledge, the highest production titers yet reported in the literature for an *E. coli* strain with the isobutanol pathway chromosomally integrated. This chromosomally integrated strain, which achieves production performance close to that of a plasmid-based strain with substantially higher pathway gene expression levels, also provides a striking example of how plasmid-based expression can be unnecessarily high. We expect that the lower expression levels in the integrants may be beneficial under more stressful conditions, alleviating unnecessary demands on the cell. Even under the relatively optimal fermentation conditions used here, the integrated strains reach higher cell densities than the plasmid-bearing strains, demonstrating the reduced burden on these strains. Furthermore, we found this approach can take advantage of beneficial gene deletions that may not be obvious choices for targeted deletion to yield optimal production.

Our results also open intriguing questions for future investigation. For example, it will be interesting to learn what factors can lead a strain with relatively high expression levels of all pathway genes, such as C1, to have such a substantially lower production level than other strains. Also intriguing is the fact that, in the highest-producing isolate we identified, C7, as well as in another high-producing isolate, B2, the gene integrations are located quite close to each other. Whether this is due to that region having optimal expression levels, or the proximity of the constructs provides some benefit, or it is merely by chance is not clear. Deep-sequencing pools of droplets collected from different degrees of screening stringency could provide further insights into whether proximity leads to improvements in production. Additionally, studies with targeted integrations could be designed to address this question.

## Materials and Methods

### Strains and plasmids

Strains and plasmids used are listed in Table S1. JCL260, pSA65, and pSA69^38, 50^ were provided by James Liao, UCLA. Keio strains were obtained from the *E. coli* genetic stock center (CGSC, http://cgsc2.biology.yale.edu/). Plasmid pSAS31 was constructed by Scott Scholz^33^.

### Media

M9IPG, consisting of M9 salts (47.8 mM Na_2_HPO_4_, 22.0 mM KH_2_PO_4_, 8.55 mM NaCl, 9.35 mM NH_4_Cl, 1 mM MgSO_4_, 0.3 mM CaCl_2_), micronutrients (2.91 nM (NH_4_)_2_MoO_4_, 401.1 nM H_3_BO_3_, 30.3 nM CoCl_2_, 9.61 nM CuSO_4_, 51.4 nM MnCl_2_, 6.1 nM ZnSO_4_, 0.01 mM FeSO_4_), thiamine HCl (3.32 μM) and dextrose (D-glucose) at the stated concentrations, was used for all culturing experiments. For SnoCAP screening, in both microdroplets and microplates, the glucose concentration was 20 g/L, and the medium was supplemented with 3 mM L-isoleucine and 50 μg/mL kanamycin. For isobutanol production monocultures, the medium contained 36 g/L glucose, 5 g/L yeast extract, and no antibiotics were added. Precultures for both screening and production cultures were carried out in LB Lennox with antibiotics appropriate to the strain. When used, antibiotics were supplied at the following concentrations: ampicillin, 100 μg/mL; kanamycin, 50 μg/mL; tetracycline, 10 μg/mL; chloramphenicol, 50 μg/mL.

### Transposon integration library construction

All primers and oligonucleotides were ordered from Integrated DNA Technologies, Inc. (IDT) and are listed in Table S2. Each of the integration constructs was amplified by PCR from the pSA69 plasmid, adding the Tn5 mosaic ends. Primers phosph_transp_kan_for and phosph_transp_alsS_rev were used to amplify *kan*-PLlacO1-*alsS* (for Libraries A, B and C). phosph_transp_PLlacO1_ilvC_for and phosph_transp_ilvD_rev were used to amplify *ilvCD* and introduce a PLlacO1 promoter in front of *ilvC* (for Library C). phosph_transp_PLlacO1_ilvD_for and phosph_transp_ilvD_rev were used to amplify *ilvD* and introduce a PLlacO1 promoter in front of *ilvD* (for Library B). The linear PCR product was then digested with both DpnI and SpeI enzymes (NEB) to digest the template plasmid, phosphorylated with T4 polynucleotide kinase (NEB), cleaned with a PCR clean-up kit (Qiagen) and eluted in TE buffer. The DNA was then reacted with EZ-Tn5 transposase (Lucigen) according to the manufacturer’s instructions. The resulting transposome was then electroporated into the appropriate strain. The cells were recovered for 1.5 hr with 1 mL SOC medium. Then 50 μL was used for dilution and plating on LB with kanamycin plates to assess library size, and the remaining cells were grown to saturation in 100 mL selective medium (either LB with 50 μg/mL kanamycin for *alsS* integration or M9IPG with 0.1 mM IPTG and 3 mM lysine for *ilvCD* or *ilvD* integration). The cells were then frozen in 1 mL aliquots (resuspended in fresh LB with 25% glycerol) and later thawed, washed, and grown in LB with 50 μg/mL kanamycin to prepare them for screening.

### SnoCAP screening

SnoCAP screening for 2-KIV production, in droplet and microplate formats, was employed as previously described^36^. Stationary phase cultures in LB were used as the inocula for both formats. For the microplate format, K12 Δ*ilvD* was used as the sensor strain. M9IPG with 20 g/L glucose, 3 mM isoleucine, and 50 μg/mL kanamycin was used as the medium for all screening co-cultures except for those with the *ilvD* and *ilvCD* libraries (without *alsS* integration), in which the kanamycin was omitted. For the droplet format, K12 Δ*ilvD* pSAS31, which expresses mNeonGreen, was used as the sensor strain. For the droplet sorting assay, the droplet collection device was soaked in a mixture of HFE-7500 oil and water for several days before use in order to improve droplet stability after collection.

### Isobutanol fermentations and analysis

Isobutanol cultures were grown similarly to the method previously described^36^. Overnight cultures in LB with antibiotics were diluted 1:100 v/v into 10 mL of M9IPG with 36 g/L glucose, 5 g/L yeast extract, and no antibiotics, in a 125-mL baffled, unvented polypropylene flasks. Cells were grown for 2.5 to 3 h at 37 °C, 250 rpm, followed by induction of the pathway genes with 0.1 mM IPTG. Flasks were then sealed with parafilm and incubated at 30 °C, 250 rpm. Samples were taken daily to measure cell density and glucose and isobutanol concentrations. Cell density was assessed by measuring the OD_600_ of 200 μL of culture, diluted into the linear range, in a VersaMax microplate reader (Molecular Devices). Glucose and isobutanol were measured by high-performance liquid chromatography (HPLC), as previously described^37^. In the case of cultures in which RNA was harvested, 0.5 mL of cells were harvested with RNAprotect Bacteria Reagent (Qiagen) 3-4 h after induction. RNA was isolated using an RNA mini kit (Qiagen) according to the manufacturer’s instructions, including on-column DNAse I digestion. An additional digestion using Turbo DNA-free Kit (Invitrogen) was performed after purification to eliminate any remaining genomic DNA. The RNA was then reverse transcribed to cDNA using MultiScribe Reverse Transcriptase (Invitrogen) and approximately 400 ng RNA per 20 μL reaction, with the following temperature profile: 25 °C, 10 min; 37 °C, 120 min, 85 °C, 5 min. A no reverse transcriptase control was carried out for each sample and checked with at least one set of primers during the real-time PCR analysis to ensure no significant residual genomic DNA. qPCR was conducted on the cDNA using SYBR Green qPCR Master Mix (Life Technologies) with 20 μL reactions. Reactions were run in 96-well plates on a 7900HT Fast Real-Time PCR Machine (Applied Biosystems), courtesy of the University of Michigan Advanced Genomics Core. The PCR program was as follows: 50 °C, 2 min; 95 °C, 10 min; 40 cycles of 95 °C for 15 s, 60 °C for 1 min. Primers used are listed in Table S2. Data were analyzed using the 2^−ΔΔ*CT*^ method^51^.

### kivD and adhA integration

A CIChE construct consisting of *kivD, adhA*, and a chloramphenicol resistance gene, *cat*, flanked by 1 kb homology regions, was previously generated and integrated into the *aslB* locus of the NV3r1 strain by λ-Red recombineering^36^. Following the method of Tyo et al.^18^, we passaged this strain into increasing concentrations of chloramphenicol and then deleted the *recA* gene by P1 transduction from donor strain BW 26,547 Δ*recA*::*kan* Lambda *recA*+ (obtained from the Coli Genetic Stock Center (CGSC)). Transductants were selected on LB plates with 50 μg/mL kanamycin and the corresponding concentration of chloramphenicol. Genomic DNA was isolated using DNEasy Blood & Tissue Kit (Qiagen), and the copy number was determined by qPCR using the same primers as Tyo et al.^18^, with one set for the *cat* gene and one for the *bioA* gene, which is used as a single copy reference gene. Absolute copy number was determined by comparison to a strain with a single copy of *cat* integrated into the *intC* locus (NV3r1 Δ*intC::yfp-cat*). To transfer this integration into the C7 strain generated in this study, we prepared P1 lysates from NV3r1 (*recA+*) with the *kivD, adhA* CIChE construct after growing on 80 μg/mL chloramphenicol and transduced it into the C7 strain, selecting on LB plates with 40 μg/mL chloramphenicol. Resulting colonies were isolation streaked twice on LB agar with 40 μg/mL chloramphenicol and 0.8 mM sodium citrate.

## General

The authors thank Scott Scholz and Peter Freddolino for valuable discussions regarding position-dependent gene expression variation and transposon footprinting, as well as Mark Burns, Lola Eniola-Adefeso, and Chris Barr for the use of equipment in their laboratories. The isobutanol strains and plasmids were originally provided by James Liao (UCLA, formerly). The CIChE construct was constructed using the pTGD plasmid provided by Keith Tyo (Northwestern U).

## Author contributions

T.S. and X.N.L. conceived and designed the study. T.S. and A.K. constructed the integration libraries. M.T.C. and K.K. developed the droplet sorting platform. T.S. and M.T.C. performed the library screening. T.S. and D.N.C. characterized the library isolates and fully integrated strains. T.S. and X.N.L. wrote the paper.

## Supplementary Materials

## Supplementary FIgures

**Figure S1.**
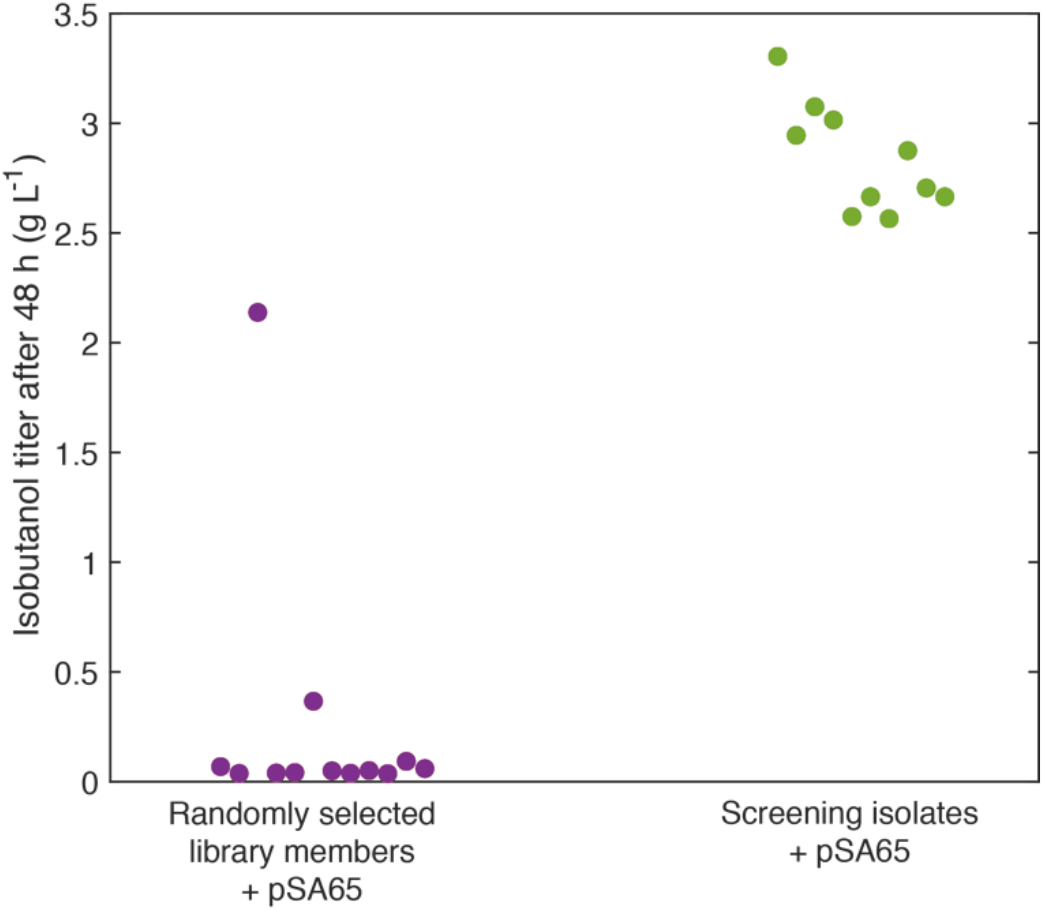
Comparison of production performance of randomly selected library members to library members isolated by screening, after transformation with pSA65. Plot shows isobutanol titers 48 h after IPTG induction of 12 randomly selected colonies from unscreened Library A (*alsS* integrated) and 10 colonies from the pool of droplets produced by screening Library A, after sorting the top 0.3% most fluorescent droplets (top 3% of library members).

**Figure S2.**
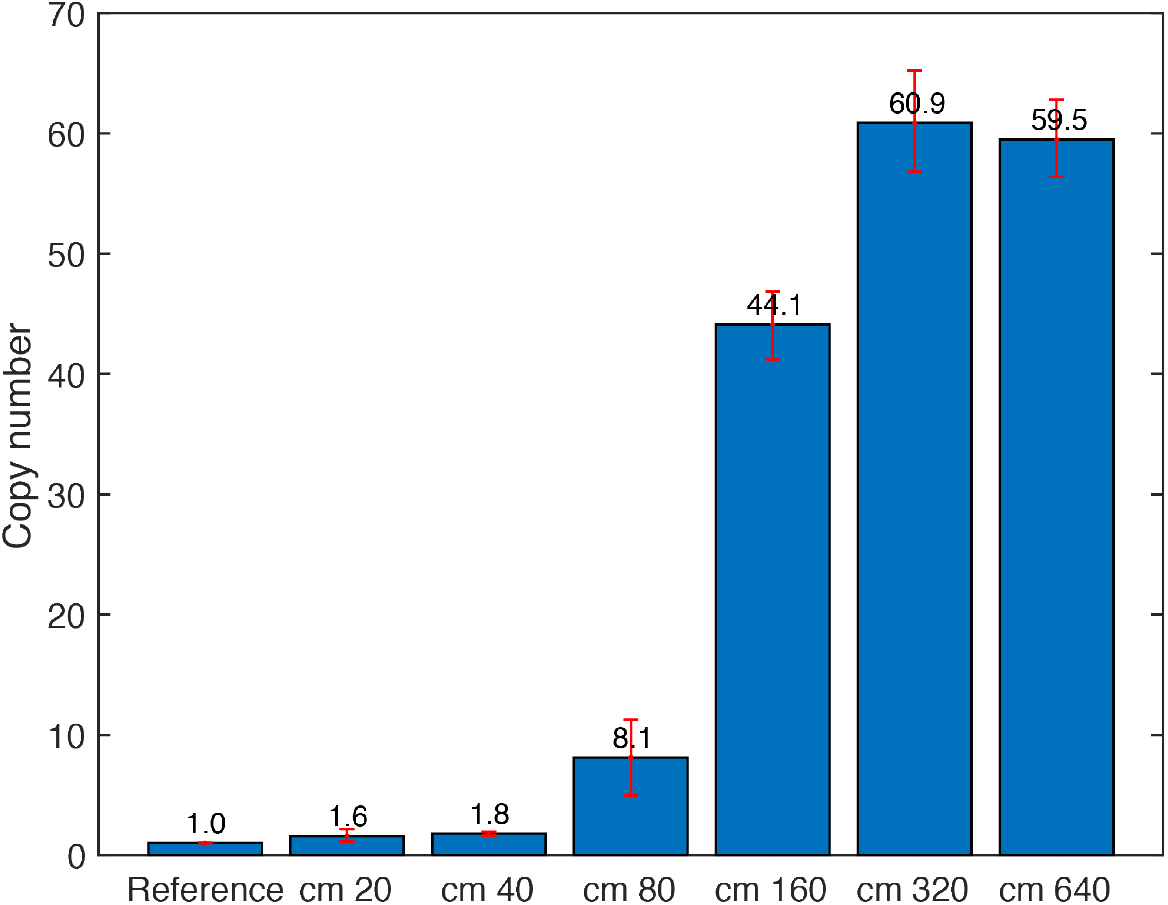
CIChE construct copy number. Copy number of a CIChE construct containing the *kivD, adhA,* and *cat* genes in the NV3r1 strain after passaging into higher concentrations of chloramphenicol and deletion of *recA*. The measurement was conducted by qPCR of the *cat* gene. “cm X” refers to NV3r1 with the CIChE construct integrated in the *aslB* site and propagated in X μg/mL of chloramphenicol before deletion of the *recA* gene. “Reference” refers to a strain with a single copy of the *cat* gene (NV3r1 Δ*intC*::*yfp-cat*). Error bars represent the standard deviation of three technical replicates.

## Supplementary Tables

**Table S1:**
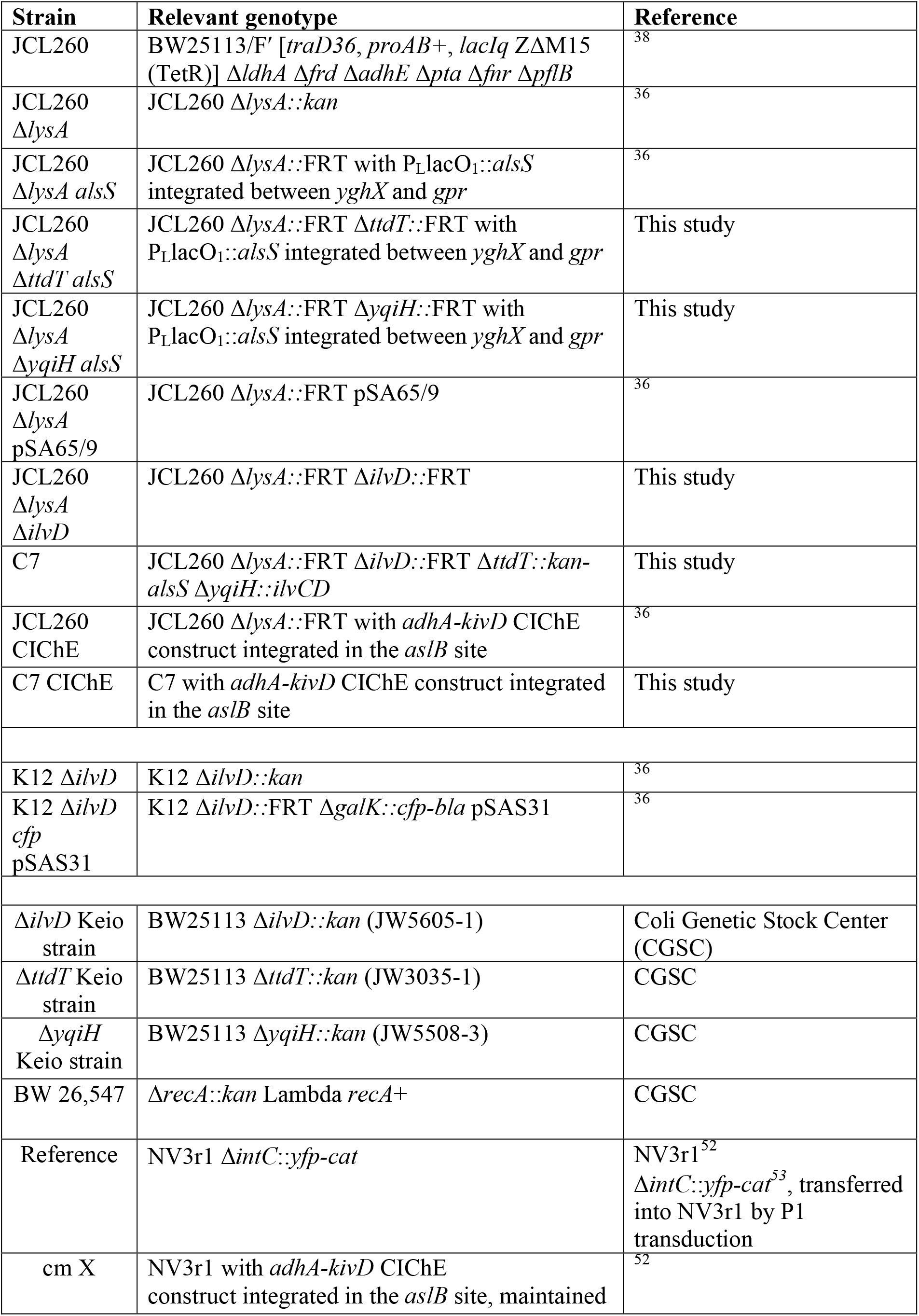
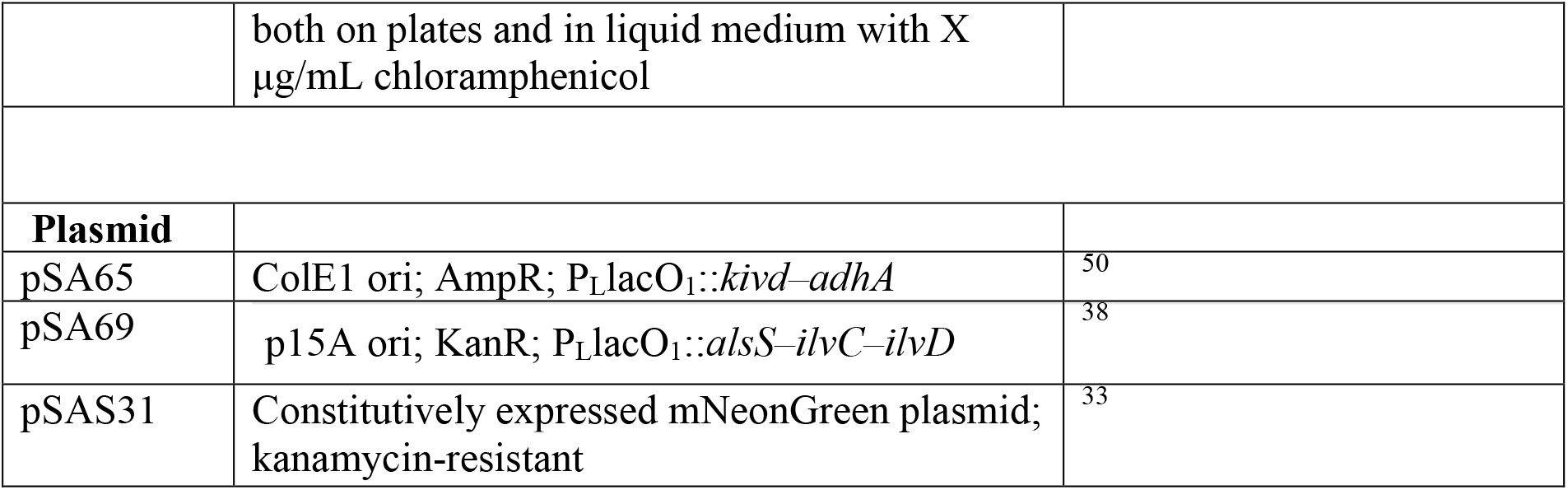
Strains and plasmids

**Table S2:**
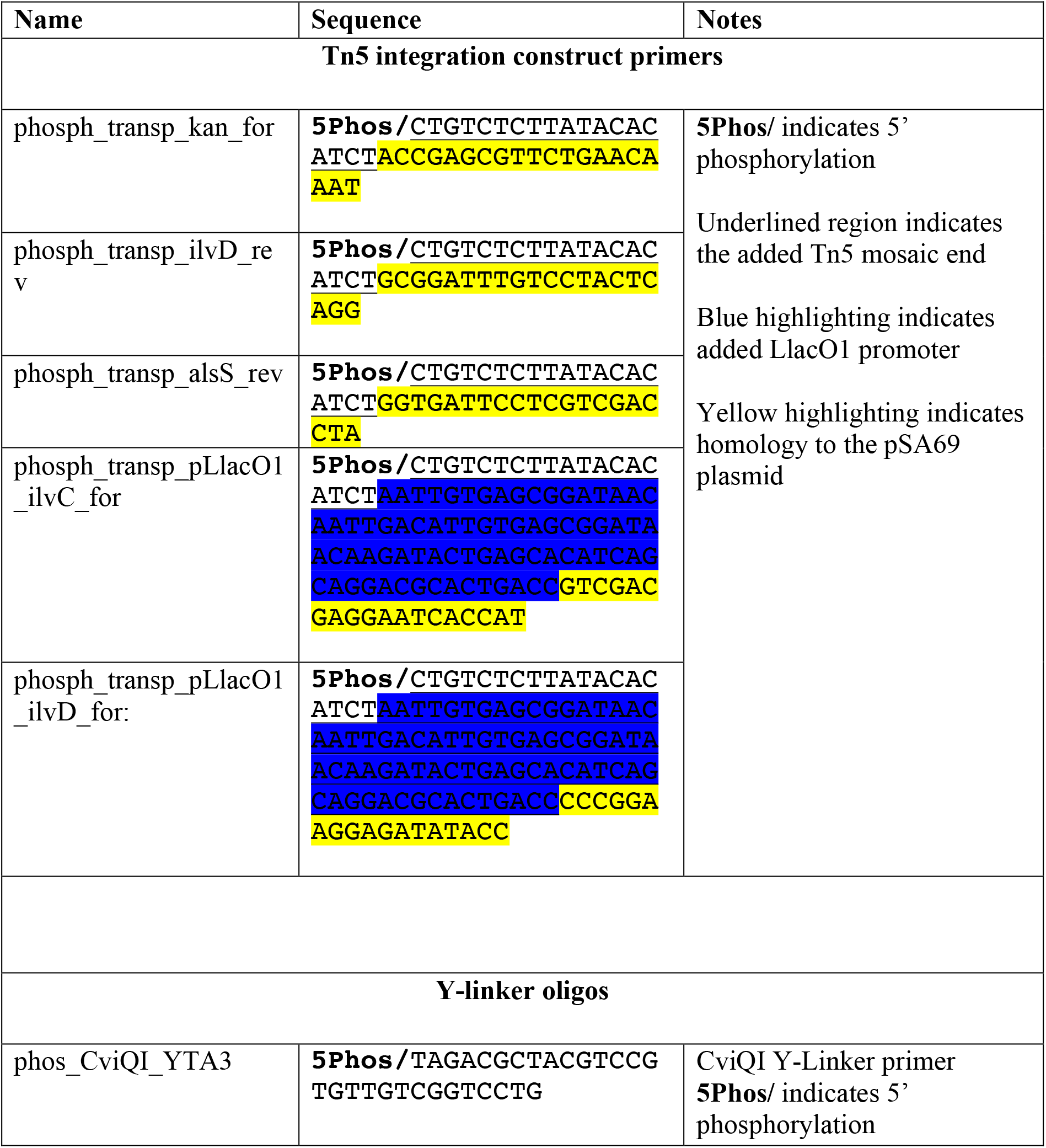
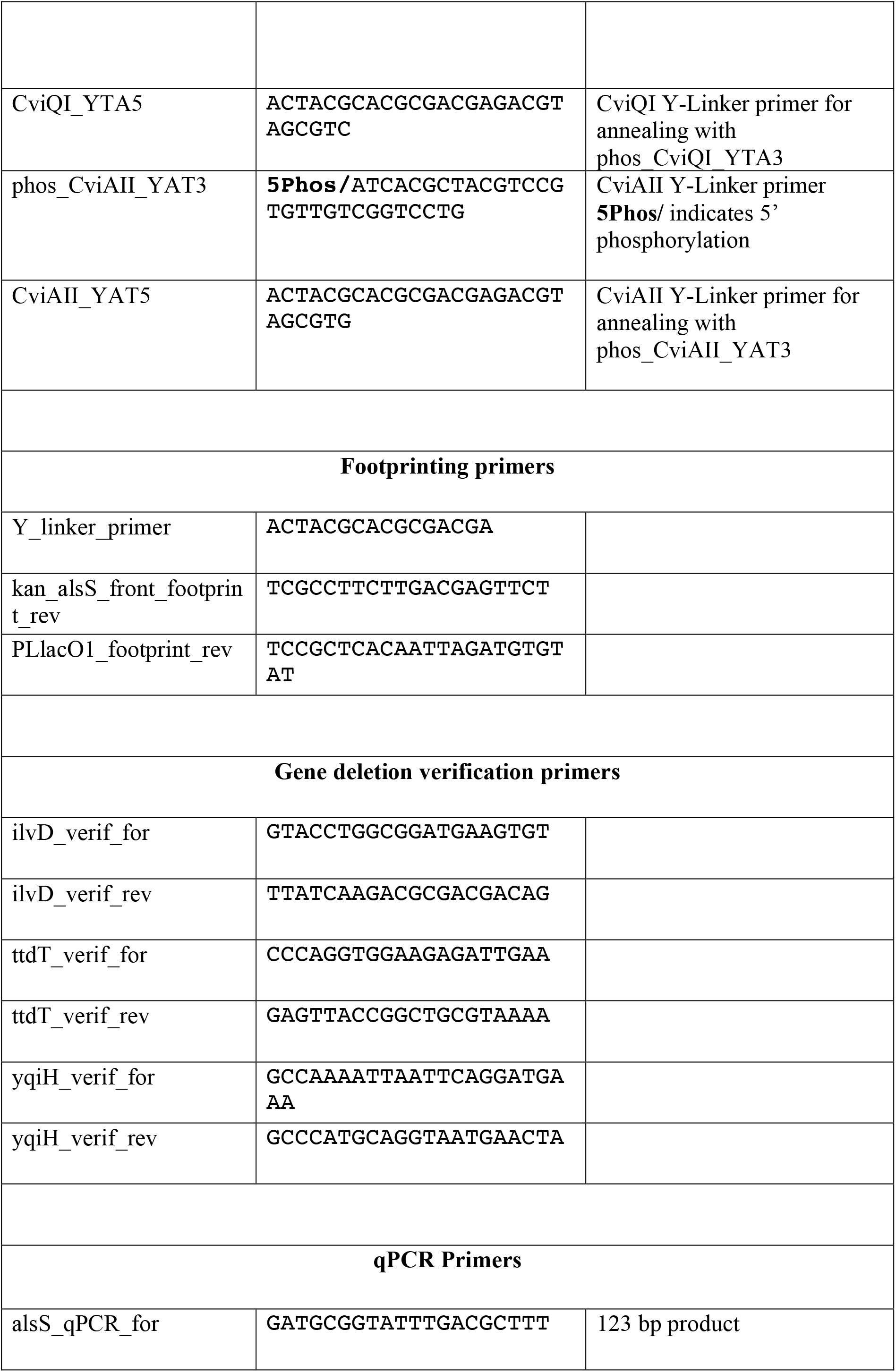
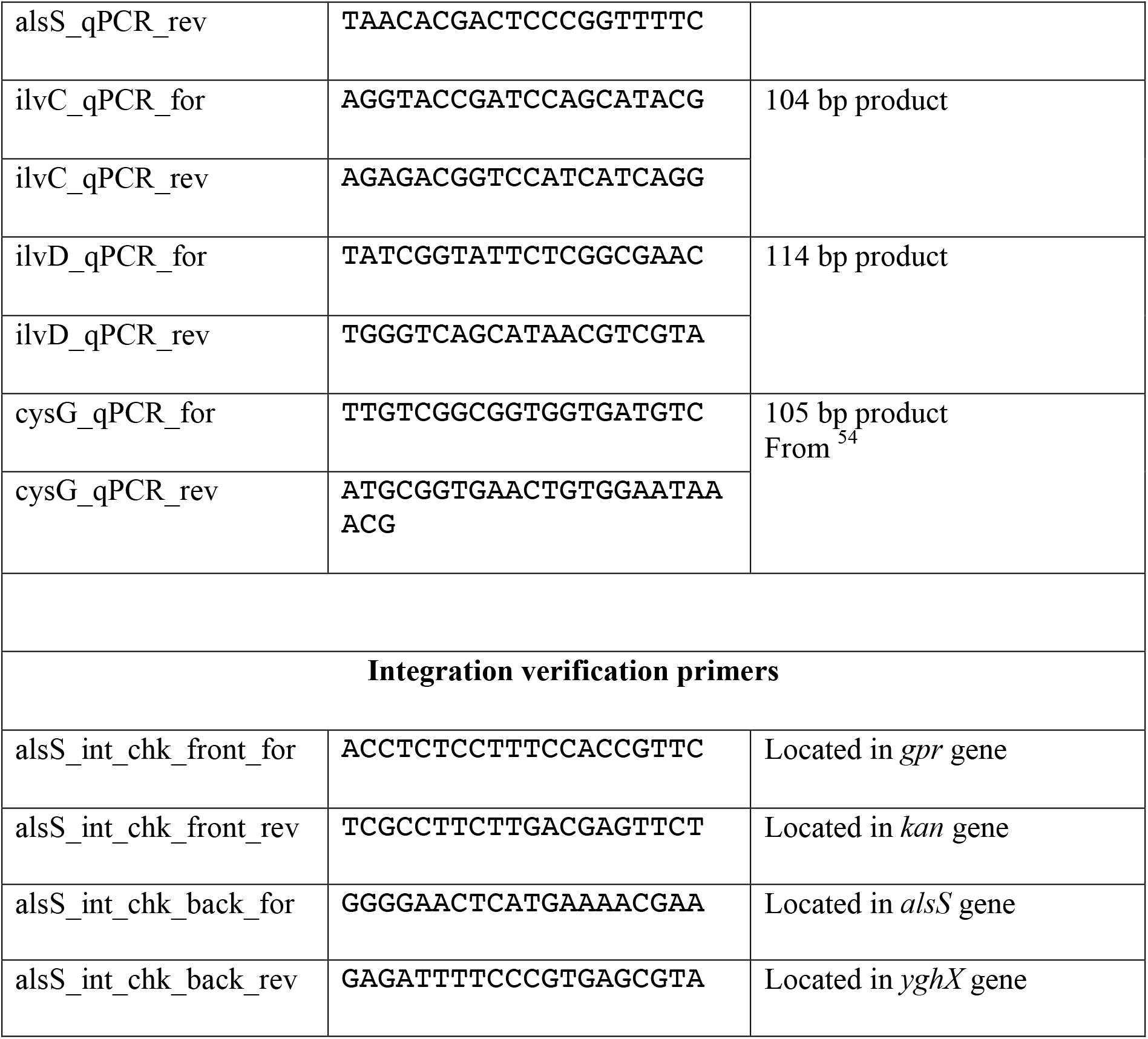
Primers and oligonucleotides

